# Predicting locus phylogenetic utility using machine learning

**DOI:** 10.1101/2024.05.06.592828

**Authors:** Alexander Knyshov, Alexandra Walling, Caitlin Guccione, Rachel Schwartz

**Author notes:** **Associate Editor:** TBA.

## Abstract

Disentangling evolutionary signal from noise in genomic datasets is essential to building phylogenies. The efficiency of current sequencing platforms and workflows has resulted in a plethora of large-scale phylogenomic datasets where, if signal is weak, it can be easily overwhelmed with non-phylogenetic signal and noise. However, the nature of the latter is not well understood. Although certain factors have been investigated and verified as impacting the accuracy of phylogenetic reconstructions, many others (as well as interactions among different factors) remain understudied. Here we use a large simulation-based dataset and machine learning to better understand the factors, and their interactions, that contribute to species tree error. We trained Random Forest regression models on the features extracted from simulated alignments under known phylogenies to predict the phylogenetic utility of the loci. Loci with the worst utility were then filtered out, resulting in an improved signal-to-noise ratio across the dataset. We investigated the relative importance of different features used by the model, as well as how they correspond to the originally simulated properties. We further used the model on several diverse empirical datasets to predict and subset the least reliable loci and re-infer the phylogenies. We measure the impacts of the subsetting on the overall topologies, difficult nodes identified in the original studies, as well as branch length distribution. Our results suggest that subsetting based on the utility predicted by the model can improve the topological accuracy of the trees and their average statistical support, and limits paralogy and its effects. Although the topology generated from the filtered datasets may not always be dramatically different from that generated from unfiltered data, the worst loci consistently yielded different topologies and worst statistical support, indicating that our protocol identified phylogenetic noise in the empirical data.

## Introduction

Unresolved phylogenetic relationships in the Tree of Life can negatively impact species classification (Som, 2015), conservation (Swenson, 2009), and our understanding of traits CITE(Avila-Lovera et al), among other topics of interest to biologists. At the same time, even modern phylogenetic analyses frequently result in a lack of resolution in some relationships, either due to poor statistical support or to conflicting results in different analyses (Chakrabarty *et al*., 2017; Esselstyn *et al*., 2017; Rodríguez-Ezpeleta *et al*., 2007; Smith *et al*., 2015). Thus, a lack of resolution in relationships is typically due to the prevalence of non-phylogenetic signal and noise.

Sources of phylogenetic error can be broadly divided into stochastic and systematic factors (Kapli *et al*., 2021). Stochastic error results from prevailing random noise in an insufficient amount of data (Kapli *et al*., 2021). However, this form of error is typically well-managed in large contemporary phylogenomic-scale datasets (Philippe *et al*., 2011). On the other hand, systematic error is due to consistent differences in the sequence data, either between taxa, between loci, or both (Kapli *et al*., 2021; Philippe *et al*., 2011). As systematic error is independent of the size of the dataset, it is generally more difficult to address (Jeffroy *et al*., 2006; Philippe *et al*., 2011). Systematic error can be caused by an array of analytical factors including data acquisition and processing artifacts (Philippe *et al*., 2011; Simion *et al*., 2017), incorrect homology inference (Fernández *et al*., 2020), alignment errors (Ranwez and Chantret, 2020), missing data (Wiens, 2006), and even software errors (Simion *et al*., 2020). While most analyses attempt to include only orthologous sequences (Boussau and Scornavacca, 2020; Fernández *et al*., 2020), the presence of paralogs is arguably one of the biggest methodological issues in modern-day phylogenomic datasets (Brown and Thomson, 2017; Simion *et al*., 2020). While in-paralog noise is minor and poses no significant issues for inference (Yan *et al*., 2022), divergent out-paralogs or deep paralogs, which predate the species split, can create significant artifactual relationships (Gatesy and Springer, 2017; Springer and Gatesy, 2018a, b). Additional systematic impacts on phylogenetic analyses can come from an array of biological properties, including a high level of Incomplete Lineage Sorting (ILS), differences in rates of molecular evolution, biases in base composition, or variation in strength of selection on the substitutions in the sequence (Degnan and Rosenberg, 2009).

One common approach to reduce systematic error is locus filtering or subsampling (Edwards, 2016; Gilbert *et al*., 2018; Smith *et al*., 2022). In this approach, loci with properties indicative of elevated noise levels are excluded from the phylogenetic inference, thus maximizing the signal-to-noise ratio (Chen *et al*., 2015; Edwards, 2016; Molloy and Warnow, 2018; Simmons *et al*., 2016). Several criteria have been used to rank loci in order to filter out those that are sub-optimal, including missing data (Brown and Thomson, 2017; Evangelista *et al*., 2021; Herrando-Moraira and Cardueae Radiations Group, 2018; Kocot *et al*., 2017; Molloy and Warnow, 2018; Mongiardino Koch and Thompson, 2021), taxon occupancy, or the presence/absence of taxa across a set of characters or traits,(Borowiec *et al*., 2015; Dietrich *et al*., 2017; Evangelista *et al*., 2021; Gernandt *et al*., 2018; Herrando-Moraira and Cardueae Radiations Group, 2018; Lemmon *et al*., 2009; Mongiardino Koch and Thompson, 2021; Young *et al*., 2016), proportion of variable or parsimony informative sites and substitution rate (Borowiec *et al*., 2015; Gernandt *et al*., 2018; Herrando-Moraira and Cardueae Radiations Group, 2018; Mongiardino Koch and Thompson, 2021; Whelan *et al*., 2015), alignment length (Brown and Thomson, 2017; Evangelista *et al*., 2021; Gernandt *et al*., 2018; Mongiardino Koch and Thompson, 2021), base composition heterogeneity (Evangelista *et al*., 2021; Kocot *et al*., 2017; Mongiardino Koch and Thompson, 2021; Whelan *et al*., 2015), saturation level (Borowiec *et al*., 2015; Herrando-Moraira and Cardueae Radiations Group, 2018; Kocot *et al*., 2017; Mongiardino Koch and Thompson, 2021), average bootstrap support of a gene tree (Borowiec *et al*., 2015; Herrando-Moraira and Cardueae Radiations Group, 2018; Mongiardino Koch and Thompson, 2021), and phylogenetic informativeness (Herrando-Moraira and Cardueae Radiations Group, 2018; Mongiardino Koch and Thompson, 2021; Townsend, 2007). Most approaches have treated filtering criteria independently and applied them sequentially, although subsampling based on a combined score of multiple different criteria has been tried as well (Herrando-Moraira and Cardueae Radiations Group, 2018).

The effectiveness of subsampling to mitigate phylogenetic noise is still debated (Molloy and Warnow, 2018; Simmons *et al*., 2016). Some authors have argued that removing data from analyses can do more harm than good (Chan *et al*., 2020). However, sequencing or initial bioinformatics is a form of subsampling that occurs by default in all datasets as no molecular datasets from diverse species can include entire genome sequences in analyses. In most cases, a reduced molecular dataset is obtained and used by selection of particular regions either in advance or after genome sequencing and processing. For example, single copy protein coding orthologs (Simão *et al*., 2015; Zhang *et al*., 2019), small conserved regions of the genome (Lemmon *et al*., 2012; Literman and Schwartz, 2021; Schwartz *et al*., 2015), or ultraconserved elements (Chakrabarty *et al*., 2017; Esselstyn *et al*., 2017; McCormack *et al*., 2012; Zhang *et al*., 2019) are frequently used in phylogenetics and are all the result of genome subsampling. Fortunately, with modern-day genome-scale data, even subsampling retains extremely large datasets with extensive information.

A caveat of subsampling is that it is only effective as a mitigation approach for particular types of non-phylogenetic signal or noise. For example, loci that experienced incomplete lineage sorting (ILS) can produce accurate gene trees that do not match the species tree. The effect of these loci is best tackled by analytical approaches such as using coalescent-aware phylogenetic software (Chifman and Kubatko, 2014; Zhang *et al*., 2018). Due to the recent emphasis on the impacts of ILS on the conflicting relationships (Edwards, 2009; Rannala *et al*., 2020), and the consequent development of tools, this type of discordance is not the focus of the present paper.

Recent work has suggested that subsampling by simultaneously analyzing several confounding factors, and ranking data based on individual or combined effects in order to filter out noisy data, can improve species tree estimation (Mongiardino Koch and Thompson, 2021). Subsampling based on high statistical support of gene trees led to the best outcomes, while ranking based on missing data was ambiguous with respect to improving phylogenetic signal ratio (Mongiardino Koch and Thompson, 2021). Loci with average substitution rates also produced the best phylogenetic inference, with rates that were too high or too low producing suboptimal results (Mongiardino Koch and Thompson, 2021). However, it is unclear whether the subsampling-induced changes in the inferred species tree were improvements, since only empirical datasets, without known true trees, were considered. At the same time, the best performing subsampling criterion was found to be Robinson-Foulds (RF) distance to the inferred species tree, a conclusion with some degree of circularity as RF distance is unknown unless the true tree is already known (Mongiardino Koch and Thompson, 2021).

In this study we examined phylogenetic data filtering by leveraging simulated datasets with controlled locus properties and known true species histories. Using machine learning in the form of a random forest regressor allowed us to predict the level of phylogenetic utility in any locus, given its properties, as well as to better understand interactions between different factors contributing to noise. Our trained model accurately predicted the phylogenetic utility of the test datasets. Reanalyzing subsets of data with high and low predicted phylogenetic utility based on the trained model provided information on the impacts of the subsetting on the topologies in empirical datasets (Fong *et al*., 2012; Liu *et al*., 2017; McGowen *et al*., 2020; Wickett *et al*., 2014). We show that multiple factors can contribute to non-phylogenetic signal, and these factors can interact. The effectiveness of model-based filtering on empirical data varied among datasets, but the subsets of loci with the most non-phylogenetic signal tended to have a different inferred topology from the known true tree and poor statistical support, suggesting these data can impact phylogenetic estimation, and removing them using this approach can benefit phylogenetic estimation.

## Results

### Phylogenetic utility of loci and their properties

We first determined how our chosen proxies for locus phylogenetic utility, namely Robinson-Foulds (RF) and weighted Robinson-Foulds (wRF) similarity between estimated gene trees and the reference simulation tree, were correlated with the properties of loci we simulated (Fig. 1). The average substitution rate, simulated as branch length, had a weak positive correlation with RF similarity (Spearman’s *ρ* = 0.153, p-value = 3.11×10^*−*208^, Fig. 1a, left). The same relationship with wRF similarity was much weaker (Spearman’s *ρ* = 0.031, p-value = 6.07×10^*−*10^, Fig. 1a, right). Locus length was positively correlated with both RF and wRF similarity, but more strongly with the former (Spearman’s *ρ* = 0.382, p-value < 2.2×10^*−*308^, Fig. 1b, left) and less strongly with the latter (Spearman’s *ρ* = 0.122, p-value = 2.34×10^*−*133^, Fig. 1b, right). When only considering paralog-free loci, the correlation with wRF similarity became stronger (Spearman’s *ρ* = 0.226, p-value = 2.18×10^*−*262^).

**FIG. 1.**
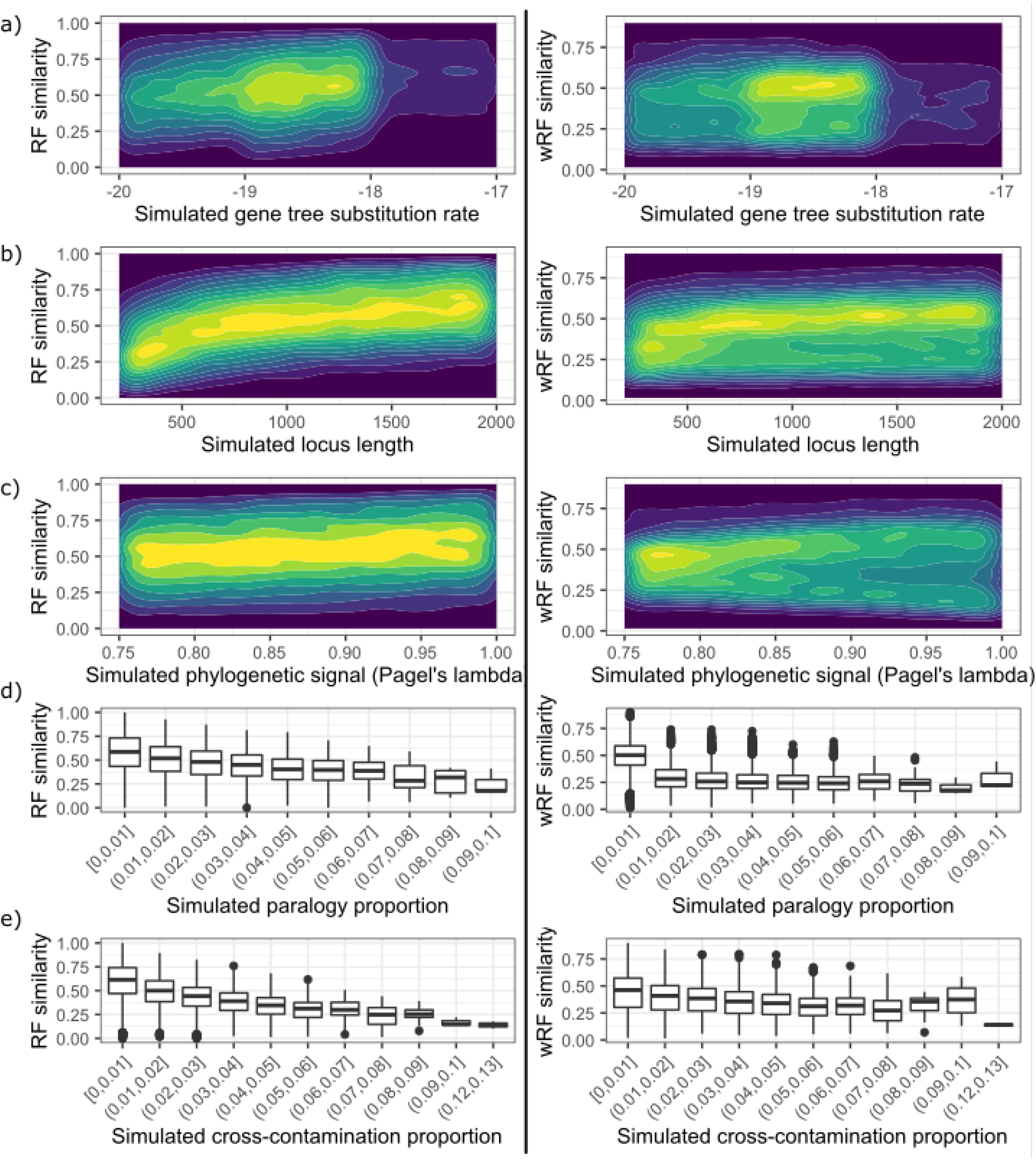
Substitution rate (panel a), locus length (b), and phylogenetic signal as represented by Pagel’s lambda (c) have weak positive correlation with the similarity of the inferred gene tree to the true species tree as measured by Robinson-Foulds (RF, left) and weighted Robinson-Foulds (wRF, right), although paralogy (d) and cross-contamination (e) have negative correlations. The association between average substitution rate and RF similarity (Spearman’s *ρ* = 0.153, p-value = 3.11×10^*−*208^) is much stronger than the association with wRF similarity (Spearman’s *ρ* = 0.031, p-value = 6.07×10^*−*10^). Locus length was positively correlated with both RF similarity (Spearman’s *ρ* = 0.382, p-value < 2.2 ×10^*−*308^) and wRF similarity (Spearman’s *ρ* = 0.122, p-value = 2.34 ×10^*−*133^), but more strongly with the former. The association between phylogenetic signal as measured by Pagel’s lambda was weakly correlated with both RF similarity and wRF similarity (Spearman’s *ρ* = 0.131 and 0.205, p-value = 1.99 ×10^*−*152^ and 4.61 ×10^*−*216^ respectively) for loci without paralogs. Due to the very high number of data points, panels a-c show 2d density plots, with yellow tones denoting high density areas where the majority of simulated loci are located while blue tones show areas of lower density. Panels d-e use box plots to better show trends in (w)RF similarity across paralogy and cross contamination bins. Because the majority of loci have near 50 values, density plots are less effective.

Phylogenetic signal, as simulated by adjusting Pagel’s lambda (Pagel, 1999) value, was weakly positively correlated with RF similarity and with wRF similarity for loci without paralogs (Spearman’s *ρ* = 0.131 and 0.205, p-value = 1.99×10^*−*152^ and 4.61×10^*−*216^ respectively, Fig. 1c). Loci with paralogs had a weak negative correlation between phylogenetic signal and wRF similarity (Spearman’s *ρ* = −0.217, p-value = 1.26×10^*−*182^), presumably as an increased proportion of internal branches amplified paralogous signal. Both RF and wRF similarity were negatively correlated with cross-contamination proportion (Spearman’s *ρ* = −0.452 and −0.222 respectively, p-values < 2.2×10^*−*308^, Fig. 1e) and proportion of “deep” paralogs (Spearman’s *ρ* = −0.286 and −0.712 respectively, p-values < 2.2×10^*−*308^, Fig. 1d). Although RF similarity had a stronger correlation with cross-contamination than wRF similarity, the latter had a stronger correlation with paralogy than the former. At the same time, correlation of RF and wRF similarity with missing taxa (reduced occupancy) was low (Spearman’s *ρ* = 0.038 and 0.044 respectively, p-value = 3.11×10^*−*14^ and 2.30×10^*−*18^ respectively). Alignment score (similarity between inferred alignment and true simulated alignment) was negatively correlated with RF similarity (Spearman’s *ρ* = −0.244, p-value < 2.2×10^*−*308^), but weakly positively correlated with wRF similarity (Spearman’s *ρ* = 0.033, p-value = 1.28×10^*−*11^). We further checked the relationship between alignment score and its impacts on gene tree inference (either RF or wRF similarity between a gene tree reconstructed from an inferred alignment vs true alignment). Results showed that RF similarity between gene trees built on inferred and true alignments was negatively correlated with alignment score (Spearman’s *ρ* = −0.344, p-value < 2.2×10^*−*308^), although similar correlation with wRF similarity was positive (Spearman’s *ρ* = 0.226, p-value < 2.2×10^*−*308^). When loci with inferred alignments nearly identical to true (score *>*= 0.99) were removed from the test, both RF and wRF similarity between gene trees became positively correlated with alignment score (Spearman’s *ρ* = 0.305 and 0.484 respectively, p-values = 1.04×10^*−*35^ and 1.49×10^*−*94^ respectively).

### Phylogenetic utility of loci and its effects on difficult nodes

To justify the use of RF similarity as a metric to optimize for phylogenetic utility, we examined the relationship between RF similarity of gene trees, accuracy, and support values. Gene trees with high RF similarity to the simulation reference also had higher median scores of the Shimodaira-Hasegawa approximate likelihood ratio test (SH-aLRT) (p-values < 2.2×10^*−*308^) and UltraFast Bootstrap (p-values <= 4.56×10^*−*163^) support, and higher median proportion of correct nodes (p-values < 2.2×10^*−*308^)

Typically, not all branches or nodes of a species tree are impacted by locus tree error to the same extent. Because we aimed to use the overall locus error as the proxy for its phylogenetic utility, we investigated how the error is distributed along the simulated species trees. For all branch length bins, the proportion of correct nodes was significantly higher in gene trees with high RF similarity to the true tree (p-values <= 4.2×10^*−*7^) (Fig. 2 left panel a). Gene trees with high RF similarity to the true tree had higher bootstrap support on all but the shortest branches. Branches shorter than 0.00489 (ln(*branch length*) = −5.32) had statistically indistinct median UFBoot values regardless of the overall locus RF similarity (p-values *>*= 0.223) (Fig. 2 left panel b). Branches shorter than 0.0000905 (ln(*branch length*) = −9.31) additionally had statistically indistinct median SH-aLRT across RF similarity bins (p-values *>*= 0.306) (Fig. 2 left panel c). For all node height bins, gene trees with high RF similarity also had more correct nodes (p-values <= 4.74× 10^*−*185^), with higher SH-aLRT (p-values <= 7.74×10^*−*7^) and UFBoot (p-values <= 0.000879) support (Fig. 2 right panels a-c).

**FIG. 2.**
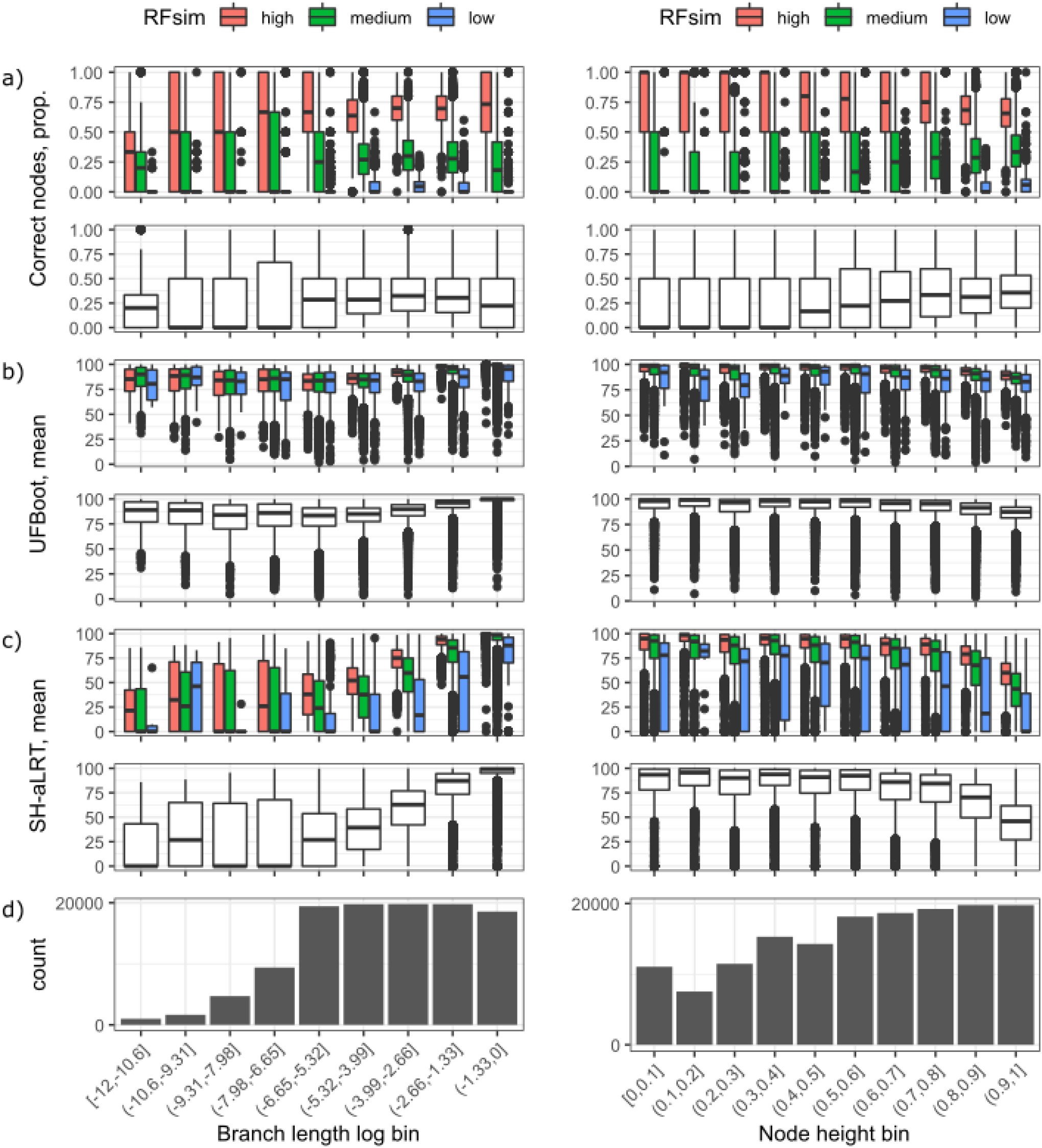
Loci with high Robinson-Foulds similarity (red) had a higher number of correctly inferred nodes and higher average statistical support. Panels show the relationship between branch length (left panels) or node hight (right panels) and the proportion of correct nodes (a) or their statistical support measured as UltraFast Bootstrap (b) and Shimodaira-Hasegawa approximate likelihood ratio test (SH-aLRT) (c). Branch lengths were binned on a ln scale to better examine short branches. Counts of nodes in each branch length or node height bin are provided for reference (d). Shorter branches (left-most branch length bins) are difficult to correctly reconstruct as seen by lower values in a-c panels, regardless of overall Robinson-Foulds similarity (indicated in red (high) green (medium) and blue (low)).

### Model training on simulated data

We trained two models, one to predict RF similarity of a locus tree to the species tree, the other to predict wRF similarity. Both models showed comparable performance (training set *R*^2^ = 0.789 and 0.84 for the RF- and wRF-based models respectively, testing set *R*^2^ = 0.616 and 0.691 respectively) (Fig. 3). The best determined values of tunable parameters for the RF / wRF model were as follows: the maximum depth of a decision tree - 1000 / 20000, the maximum number of features to consider when looking for the best split - 50% / all (auto), the minimum number of samples required to be at a leaf node - 5 / 5, the number of decision trees in the forest - 5000 / 5000, the minimum number of samples required to split an internal node - 2 / 2.

**FIG. 3.**
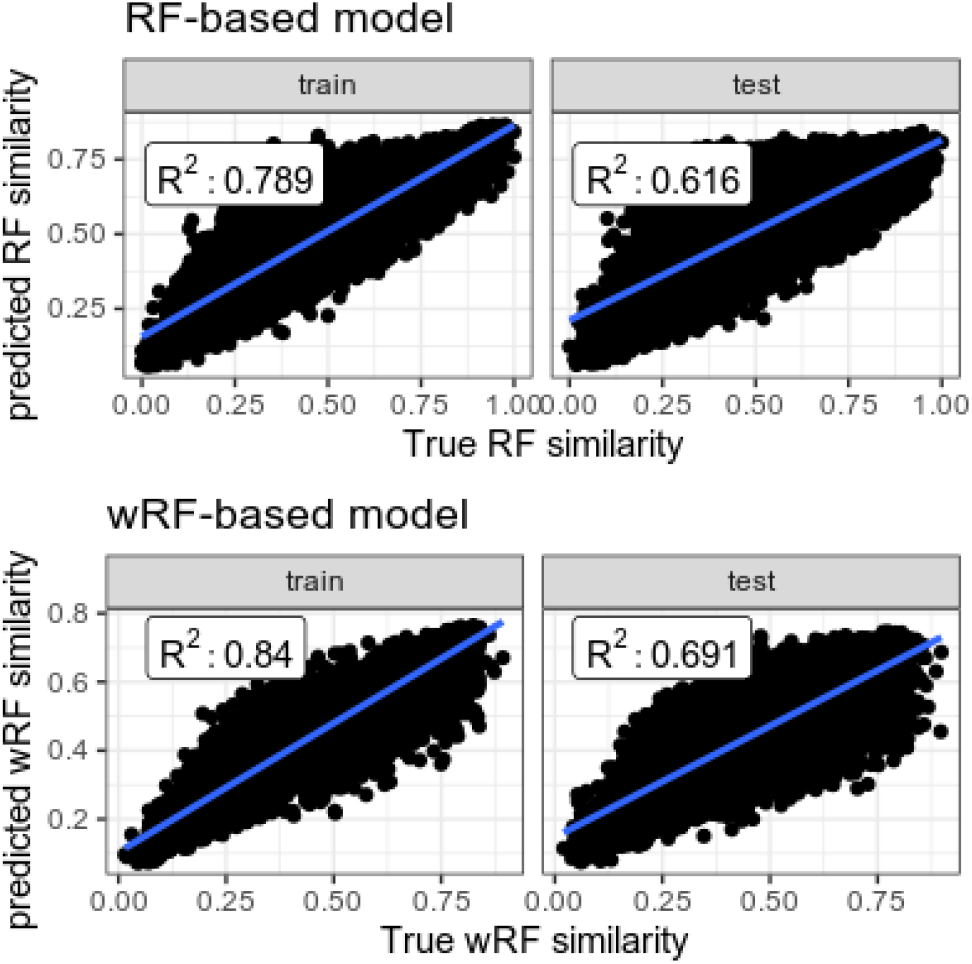
Training results of the RF-based model (top panels) and wRF-based model (bottom panels). Scatter plots show the relationship between true and predicted values for both training and testing datasets, demonstrating a high proportion of variance is explained by both models.

Average bootstrap support was the feature with the highest permutation importance (0.97 mean permutation importance) (Fig. 4, left panel). The wRF model had a more diverse distribution of feature importances, with tree rate, tree rate variation, treeness, and average support having over 0.1 mean permutation importance (0.78, 1.24, 0.19, and 0.17 respectively) (Fig. 4, right panel). At the same time, some features, including percent variable and missing data, taxon occupancy, compositional heterogeneity, and alignment length, had near zero importance in both models (Fig. 4).

**FIG. 4.**
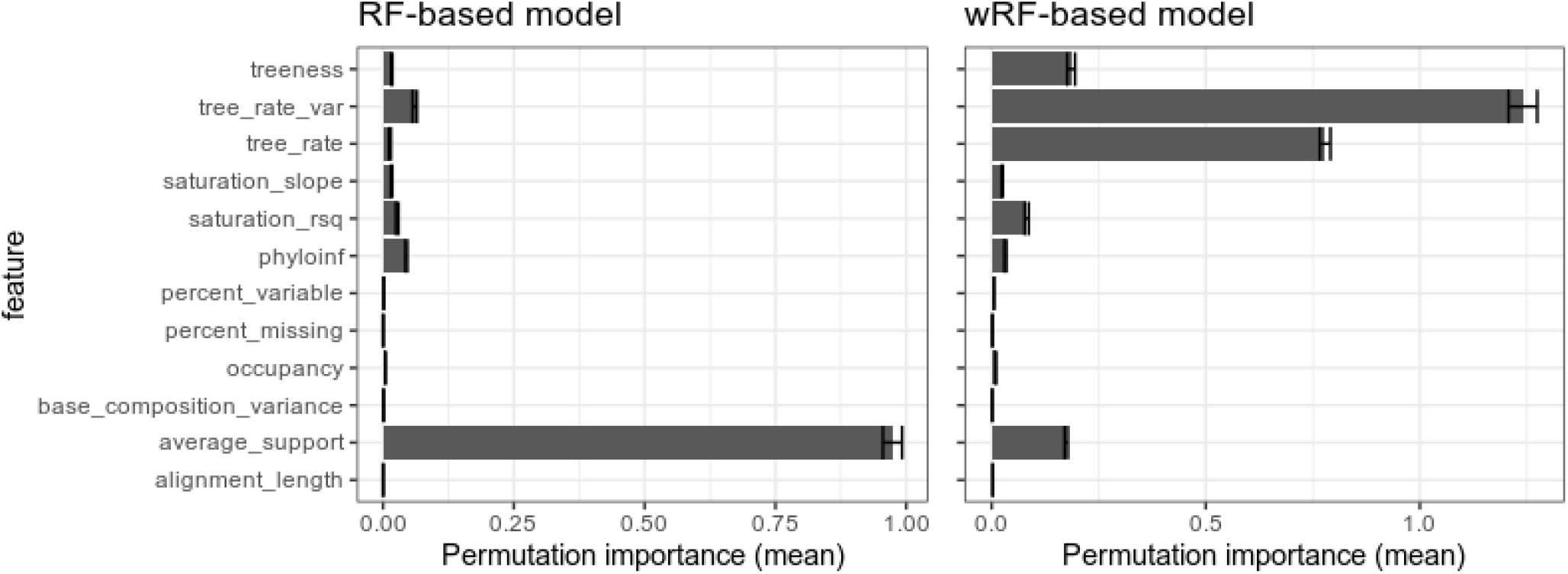
Permutation importance of each feature used in training, measuring the increase in the prediction error of the model after the feature’s values were permuted. Larger values represent larger error. For the RF model, average bootstrap support was the dominant feature, although for the wRF model several features had high importance, particularly tree-based substitution rate variance and substitution rate.

### Feature interactions

To assess the level of feature interactions in our model, we computed first-order Friedman’s H-statistic, which shows the overall level of interaction of a given feature with others defined as the share of variance explained by the interaction, and second-order Friedman’s H-statistic, which shows the level of interactions between specific features.

Tree-based substitution rate variation, average bootstrap support, and phylogenetic informativeness had the highest degree of interaction in the RF model, with 17-28% of the variance explained per feature by the interaction (Fig. 5, top left panel). Among the top most interacting feature combinations were tree-based substitution rate variance and phylogenetic informativeness, tree-based substitution rate and alignment length, tree-based substitution rate and tree-based substitution rate variance, and alignment length and percent missing data (Fig. 5, top right panel). For example, in the interaction between tree-based substitution rate variance and phylogenetic informativeness, high levels of phylogenetic informativeness led to high phylogenetic utility when the tree-based substitution rate variance was low (< 0.005); when the latter was high, even maximal phylogenetic informativeness resulted in smaller phylogenetic utility; when phylogenetic utility was low (< 200), even low tree-based substitution rate variance did not improve phylogenetic utility (Fig. 5, bottom left panel).

**FIG. 5.**
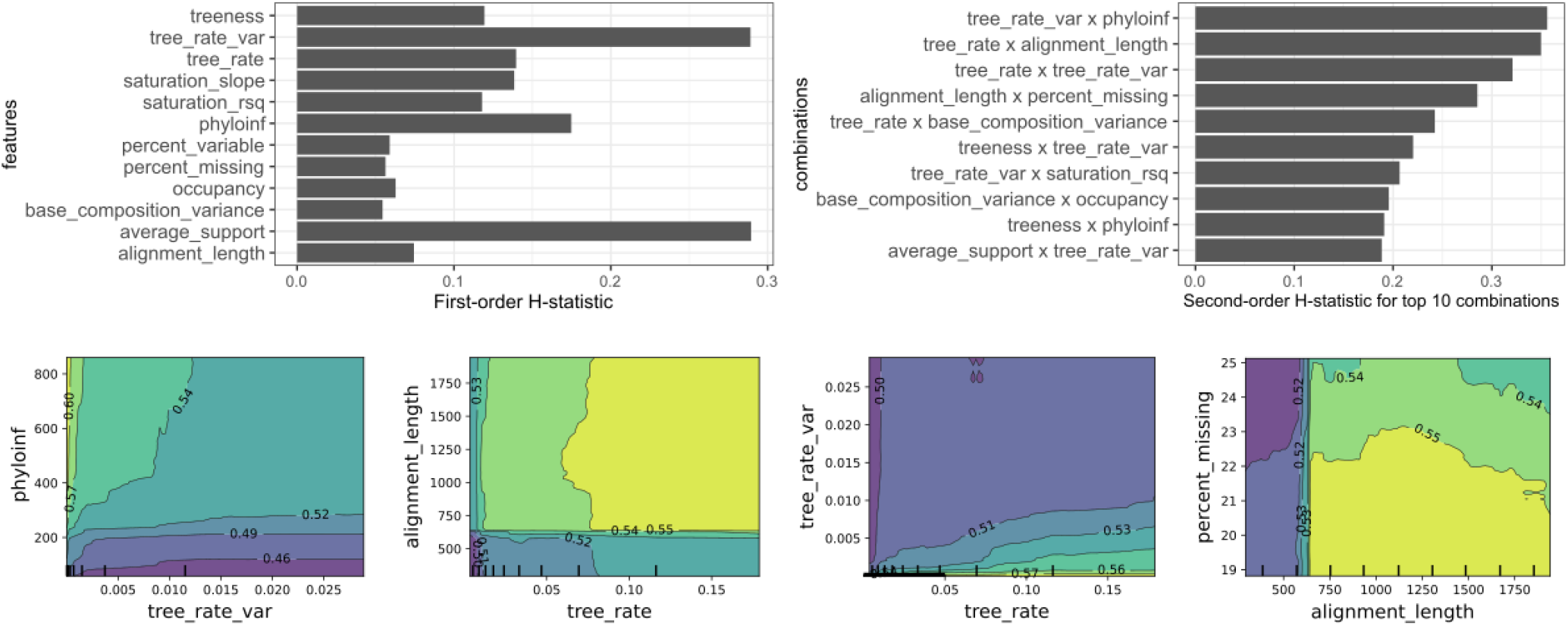
Analysis of feature interactions in the RF model. First-order H-statistic values for each feature indicate tree-based substitution rate variance and average support have the highest degrees of interaction. The top right panel shows the top ten pairwise interactions based on the second-order H-statistic values, with the strongest interaction detected between tree-based substitution rate variance and phylogenetic informativeness. Bottom panels show partial dependency plots for the top four pairwise interactions, demonstrating non-linearity in the interactions. Lines and colors delimit areas with different locus utility, from low (purple shades) to high (yellow shades).

Tree rate variation, tree rate, and average support had the highest degree of interaction in the wRF model (Fig. 6, top left panel). Among the top most interacting feature combinations were tree rate and tree rate variation, average support and tree rate variation, treeness and saturation slope, tree rate variation and saturation regression *R*^2^ (Fig. 6, top right panel). Interestingly, the tree rate and tree rate variation combination had a high level of interaction in both models (3rd highest in RF model, Fig. 5, second right bottom panel, and 1st in wRF model, Fig. 6, left most bottom panel), with high tree rate and low tree rate variation leading to high phylogenetic utility in both models, although the landscape of interaction differed between models.

**FIG. 6.**
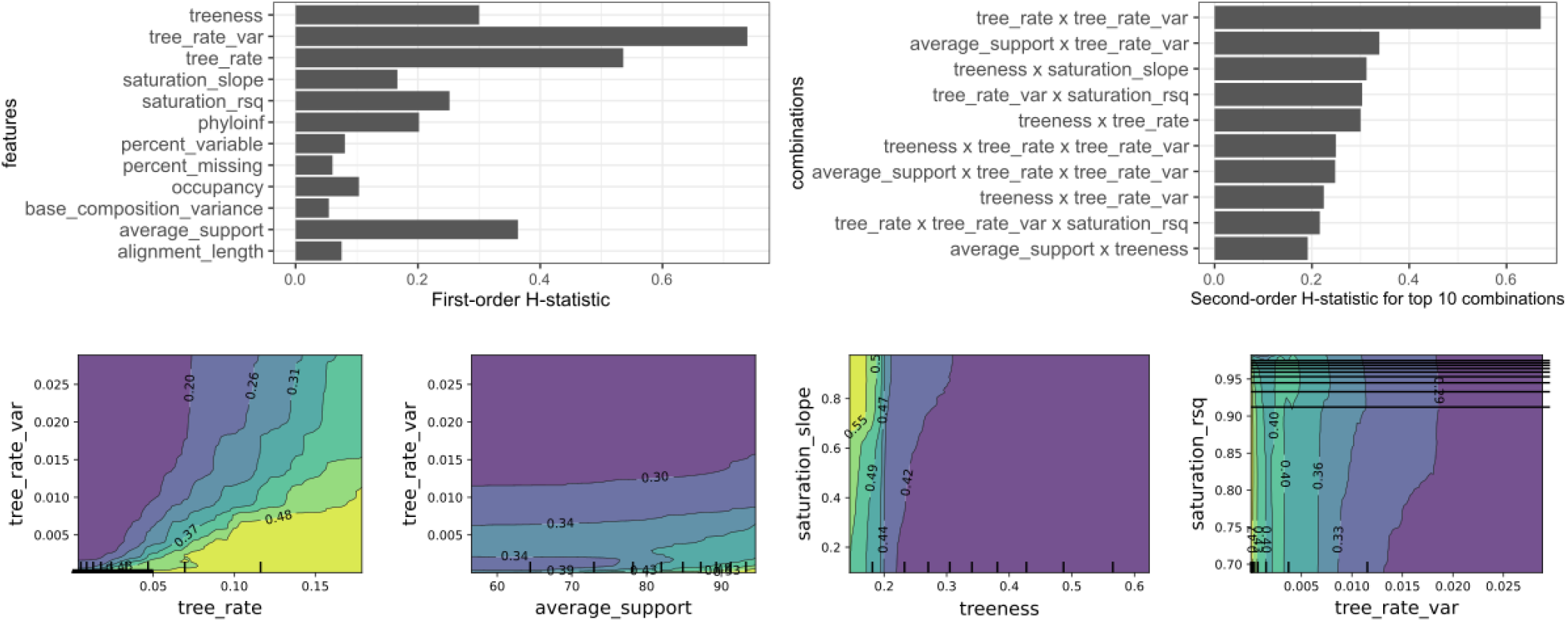
Analysis of feature interactions in the wRF model. First-order H-statistic values for each feature indicate tree-based substitution rate variance, tree-based substitution rate, average support, and treeness all have very high degrees of interaction. The top right panel shows the top ten pairwise interactions based on the second-order H-statistic values, with the strongest interaction detected between tree rate and tree rate variation. Bottom panels show partial dependency plots for the top four pairwise interactions, demonstrating non-linearity in the interactions. Lines and colors delimit areas with different locus utility, from low (blue shades) to high (yellow shades).

### Model-based subsampling of simulated loci and phylogenetic reanalysis

To examine the impact of filtering based on model predictions, we analyzed several subsets of loci (all 1000, randomly selected 600, predicted best 600, predicted best 800, and predicted worst 600) for both the Robinson-Foulds and weighted Robinson-Foulds models. The worst 600 loci subsets in both the RF and wRF models had lower median values of RF similarity in concatenation (RF: 0.932 and wRF: 0.937) and coalescence (RF: 0.976 and wRF: 0.985), and wRF similarity in concatenation (0.532 and 0.543), than those values in the best 600 loci subsets (concatenation: 0.958, 0.964, coalescence: 1, 0.996, wRF concatenation: 0.584, and 0.598 respectively). The RF and wRF values for the worst 600 loci subsets were also lower than those in the set containing all simulated loci (0.971, 0.996, 0.570) or random set (0.95,0.99,0.571).

### Progressive model-based subsampling and phylogenetic reanalysis of empirical datasets

After verifying performance of our trained models on the simulated datasets, we used the model to predict phylogenetic utility of the four empirical datasets [“Fong” (Fong *et al*., 2012), “Wickett” (Wickett *et al*., 2014), “McGowen” (McGowen *et al*., 2020), and “Liu” (Liu *et al*., 2017)]. Following the same methodology as for the simulated datasets, we assessed properties of empirical loci, predicted their utility, ranked loci, and created several subsets of data with progressively fewer loci with predicted non-phylogenetic signal. We also created subsets of the best and worst loci to examine the extreme cases. We first examined the overall differences among topologies, as shown by PCoA analyses. The 10% worst loci subsets had a consistently different topology from the rest of the subsets (Fig. 7 bottom panels). The topologies generated from the 10% best loci subsets were different from those generated by most of the other datasets, however the degree of topological difference was typically smaller (Fig. 7 bottom panels).

**FIG. 7.**
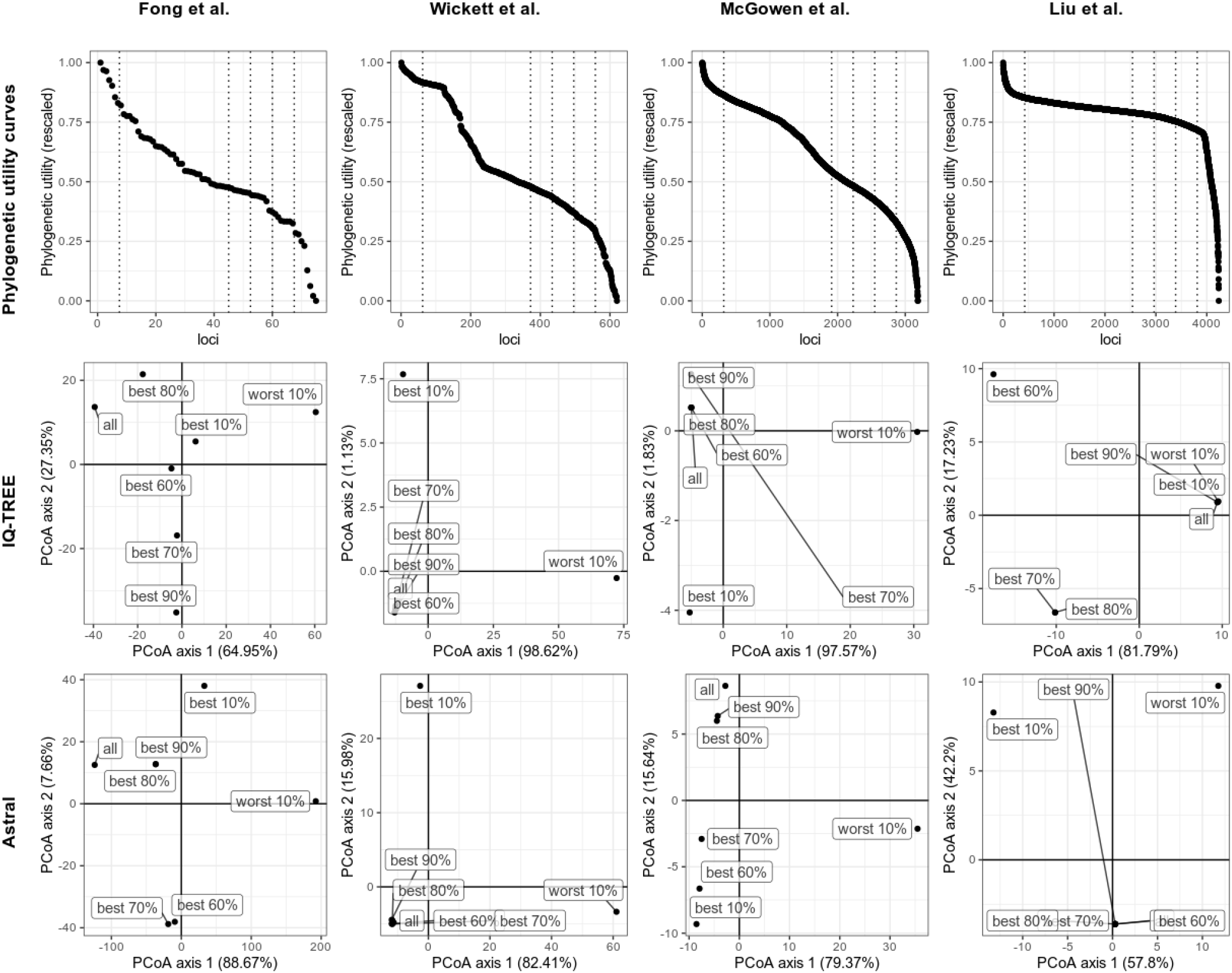
Ranking of loci in order of predicted utility for each of the four empirical datasets showing where subset cutoffs were made (top row). Results of the principle coordinate analysis (PCoA) of distances between all and filtered subsets (bottom two rows). Only the first two PCoA axes are shown. The x-axis explains the majority of the variance (57.8%-98.62%) and shows the most spread between the worst 10% subset and other filtering options, consistent with the worst subset containing maximal phylogenetic noise. The distance between the tree from the data predicted to have the most utility and other trees may be due to the size and therefore limited information in these datasets.

**FIG. 8.**
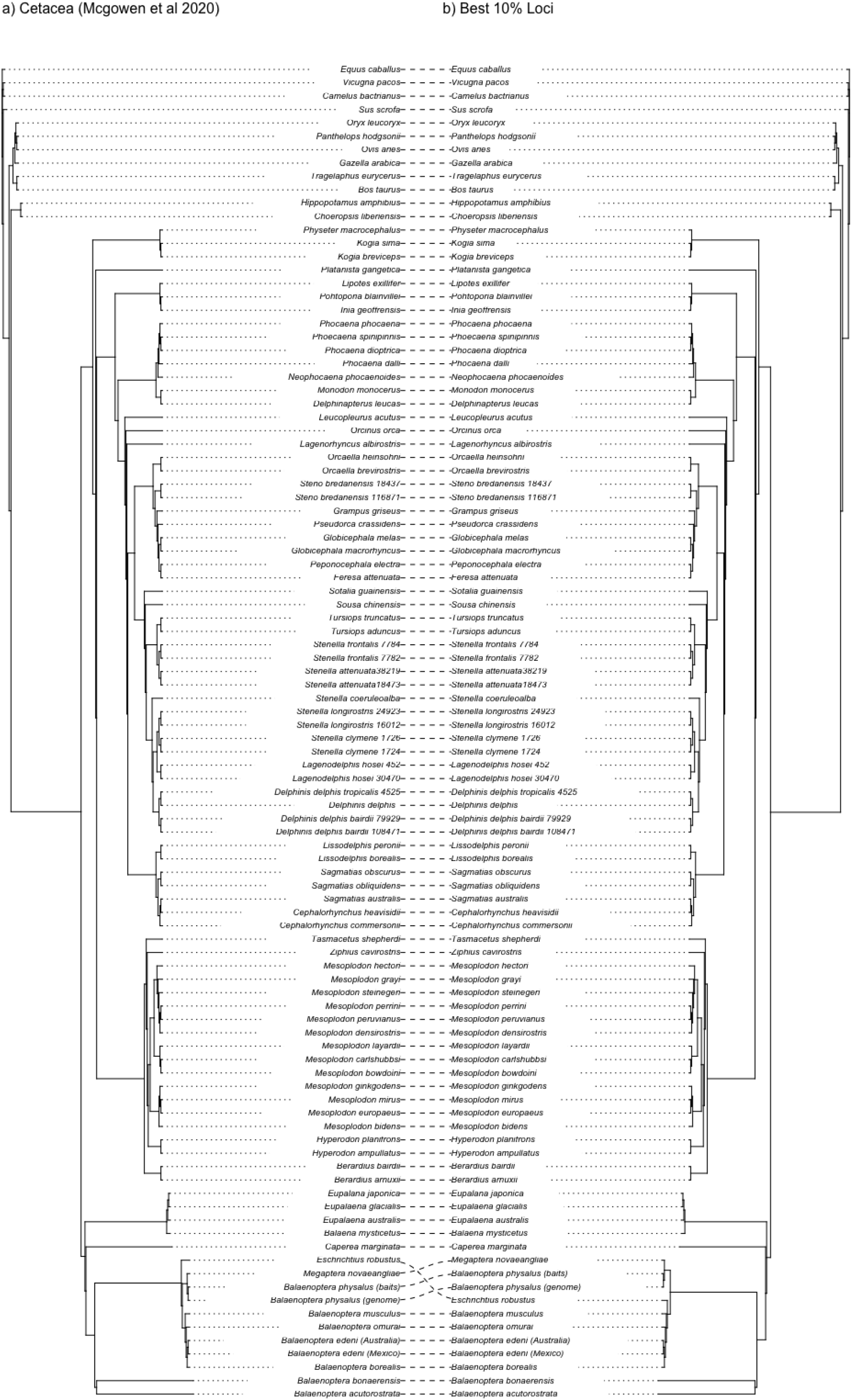
Example tanglegram comparing true species tree inferred from McGowen et al (2020) loci and the phylogeny produced from the best 10% of filtered empirical loci. While overall topologies between the two trees are congruent, the position of *Eschrichtus robustus* is rearranged within the clade corresponding to Balaenopteridae.

We then examined certain relationships in each empirical dataset that were either focal in the original study or shown to be problematic in subsequent investigations. In the Fong dataset, all loci in concatenation yielded the topology with turtles sister to crocodilians with 84.3% SH-aLRT and 55% UFBoot support. Progressive filtering using either of the model predictions did not change that topology. However, it did improve the support of that placement of turtles to 92.5% SH-aLRT and 95% UFBoot (top 70% loci using the RF model predictions) and 99.2% SH-aLRT and 93% UFboot (top 60% loci using wRF model predictions). The top 10% best loci subsets also favored the crocodilians-sister hypothesis, with comparably strong support in both RF and wRF subsets (93.5% / 99% and 97% / 99% respectively). On the contrary, the 10% worst loci subsets inferred different relationships in both models, with the worst loci as inferred by the RF model reconstructing Squamata sister (85.3% / 94%) and the worst loci as inferred by the wRF model yielding Archosauria sister (83.9% / 90%). Additionally, several taxa had incorrect position and / or excessively long branches (Accipiter placed outside Aves on a long branch, Leucocephalon and Rana on a long branch). Progressive filtering eventually mitigated these artefacts (all were gone in the RF-based 60% best subset, and all except Leucocephalon were gone in the wRF-based 60% best subset). Both top 10% best subsets contained none of these artefacts.

For the Liu dataset filtering, we focused on the relationships highlighted by Gatesy and Springer (2017). However, our ML analysis of all loci found only one of these problematic relationships, namely non-monophyletic Odontoceti, indicating that some of the Liu *et al*. (2017) controversial results might have been caused by specific phylogenetic analysis methodology of the original study. Progressive filtering did not alter the one remaining problematic relationship and it persisted in almost all subsets with the maximal support, although the RF-based 10% best loci subset had a lower support for this topology (87.6% / 92%).

Filtering of the Wickett dataset did not produce substantial changes to topology. However, the 10% worst loci selected by both models resulted in a change of topology where Hornworts were recovered as sister to Mosses and Liverworts.

For the McGowen dataset filtering, filtering of the best 10% of loci changes the placement of *Eschrichtus robustus* within Balaenopteridae, while filtering for the worst 10% of loci recovers the same placement of Eschrichtus robustus as in the “true” species tree but results in an altered topology for *Stenella coeruleoalba* within Delphinidae.

## Discussion

We trained two models to identify loci with high phylogenetic utility across a variety of simulated and empirical datasets. Our models successfully predicted loci whose gene trees had greater or lesser similarity to the true species tree as measured by Robinson-Foulds or weighted Robinson-Foulds distance. While individual characteristics of loci were predictive of phylogenetic utility as suggested in prior work (Mongiardino Koch and Thompson, 2021) and demonstrated in (Fig. 1), trained models indicated that the interactions between locus characteristics are also important in models’ predicting utility. Filtering phylogenetic datasets based on these models provided improved species tree estimates in some (although not all) cases. This work holds promise for future analysis of phylogenetic data and distinguishing among competing phylogenetic hypotheses.

### Gene tree similarity to the true species tree can be predicted using signatures in data

Importantly for this work, (w)RF distance appears to be a good proxy to optimize when filtering phylogenetic data to improve species tree estimation. We demonstrated that loci with high (w)RF similarity had a higher number of correctly inferred nodes and higher average statistical support for our simulated loci, indicating that (w)RF similarity is a good proxy for gene tree error.

The random forest regression models were both highly predictive of RF and wRF similarity to the true tree, with comparable performances on training and test datasets of simulated loci. Analysis of feature importance identified different features as being most predictive of (w)RF similarity between the RF and wRF models. For the RF model, first-order H-statistic values (Friedman and Popescu, 2008) for each feature indicate tree-based substitution rate variance and average support have the highest degrees of interaction; the strongest interaction detected was between tree-based substitution rate variance and phylogenetic informativeness. For the wRF mdoel, however, average support was less important than tree-based substitution rate variance, with treeness and tree rate also important; the strongest interactions detected were between tree-based substitution rate variance and phylogenetic informativeness, followed by tree-based substitution rate variance and alignment length.

When the (w)RF models were applied to filter the best and worst subsets of simulated loci for downstream phylogenetic inference, the 600 loci predicted to be least useful produced estimated species trees more different from the true tree than the loci predicted to be have the most utility. The worst loci subset also produced lower support values, as would be expected due to conflicting and erroneous signal.

### Model properties

Analysis of feature importance confirms some factors found by Mongiardino Koch and Thompson (2021): longer loci produced trees with higher similarity to the true species tree. However, Mongiardino Koch and Thompson (2021) also predicted better loci would have greater cross-contamination and with metrics of greater signal, which we did not observe. Our finding that average (bootstrap) support of a locus gene tree is a significant predictor of the phylogenetic utility of a locus also confirms the conclusions of Mongiardino Koch and Thompson (2021). Similarly, our finding that substitution rate only weakly correlates with utility is similar to what was shown in Mongiardino Koch and Thompson (2021), where average rates provided best signal.

### Filtering data

Filtering based on the model-predicted RF similarity to the true tree improves the statistical support of reconstructed relationships. This is consistent with the observation that incorrect hypotheses are highly supported in trees with low RF similarity. Additionally, filtering based on RF similarity should increase likelihood of inferring correct splits and obtaining higher statistical support at all node heights, both near the root and near the tree terminals.

While short branches remained difficult to reconstruct (filtering can remove noise but not add signal), loci whose gene trees had high RF similarity to the true species tree had a greater proportion of correct nodes; thus predicting these loci remains a positive step. For these branches bootstrap support was consistently high, even for low similarity loci, suggesting that filtering loci with low predicted RF similarity could remove support for erroneous alternate hypotheses. Unfortunately, for very short branches, the gains from using loci with higher phylogenetic signal were minor. Thus, this model-based filtering may not improve results in cases of rapid species diversification.

### Applicability of models to empirical data

When we applied our models to the empirical datasets upon which our simulations were based (Fong *et al*., 2012; Liu *et al*., 2017; McGowen *et al*., 2020; Wickett *et al*., 2014), we found that the precise level of data to include and exclude while subsampling can be challenging and somewhat arbitrary. Phylogenetic utility curves for loci indicate drop-offs in locus quality can differ from dataset to dataset; for this reason, we created multiple subsets of locus data for each empirical dataset to test performances of the best and worst loci. Principle components analysis for each of the four empirical datasets shows the greatest separation between the worst 10% of loci and all others; however, we found that the best 10% subsets were not ideal for phylogenetic inference. Filtering too drastically may bring stochastic noise into play (Chan *et al*., 2020). It is likely, that while top 10% loci may have a superior utility, they could have increased impacts of stochastic noise due to the small size of those subsets. We speculate that the shape of the phylogenetic utility curve could be examined, with drastic changes in the shape chosen for subsampling boundaries. In cases where the curve is fairly linear, an arbitrary cutoff in the range of 5-40% could be warranted, while balancing the amount of data and stochastic error.

Following loci filtering, phylogenetic analysis of the four empirical datasets provided some insight into ongoing questions in the evolutionary histories of tetrapods (Fong *et al*., 2012), mammals Liu *et al*. (2017), Cetacea McGowen *et al*. (2020), and land plants (Wickett *et al*., 2014). Analysis of the “Fong” dataset indicated that progressive filtering of loci did not change the topology with regards to the placement of turtles sister to crocodilians, but did improve support values, while the worst 10% of loci inferred different relationships for Squamata and Archosauria in both the RF and wRF models. In the “Liu” dataset, progressive filtering did not alter the problematic relationship of non-monophyletic Odontoceti, although the RF-based best 10% of loci had a lower support for this topology.

For the McGowen dataset, filtering with both the best 10% and worst 10% of loci changes the placement of some taxa. In the “Wickett” plant dataset, progressive filtering for the best loci did not change topologies, but the worst 10% o loci selected by both the RF and wRF models resulted in a change of topology where Hornworts were recovered as sister to Mosses and Liverworts, supporting the idea that use of the noisiest data will result in worse phylogenetic inference.

### Conclusion

Our models provide methods for subsampling loci while accounting for diverse features of the dataset. When applied to simulated loci, filtration based on model-predicted best and worst loci improved the accuracy of phylogenetic inference. At the same time, we show that even gene trees with overall good (w)RF measure have low confidence for short branches; our model-based subsampling might not be very helpful for the rapid divergences.

Performance of our models on simulated and empirical datasets may be partially dependent on the specific methods used to simulate data and introduce features such as missing data, which was introduced randomly across simulated loci rather than modeled off of properties of missing data in specific empirical datasets. Our models consider properties on a locus basis rather than relative to the overall dataset distribution; future studies could examine how to combine our approach with ways to take into account the overall dataset property distributions (de Vienne *et al*., 2012).

Application to four empirical datasets suggested small but meaningful changes in topology and support for nodes of interest. Retraining models on simulated datasets with parameters closely tied to a particular empirical dataset, rather than generalized from multiple empirical datasets, may result in improved predictions for specific empirical data.

## Material and methods

### Simulation

To create datasets to evaluate how locus parameters affect accuracy we used 4 empirical and 16 simulated species trees. The four empirical datasets were examined to obtain realistic parameter values for the simulations. These datasets (Fong *et al*., 2012; Liu *et al*., 2017; McGowen *et al*., 2020; Wickett *et al*., 2014) included a variety of taxa and examined contentious relationships. The Fong *et al*. (2012) dataset was concerned with the placement of turtles in the tree of life and included a diverse sample of taxa across tetrapods. The Wickett *et al*. (2014) dataset was used to investigate the phylogeny of land plants and had a diverse sample of all plant taxa and algal outgroups. The McGowen *et al*. (2020) dataset was focused on resolving the cetacean tree of life. The Liu *et al*. (2017) dataset was aimed at reconstructing the mammalian diversification.

All four empirical studies reconstructed the species trees; however, different methods were used in each case, and in some cases resulting tree files were not published. For consistency, the species trees were inferred de novo based on published alignments (Brown and Thomson, 2017; Fong *et al*., 2012; Liu *et al*., 2017; McGowen *et al*., 2020; Smith *et al*., 2015; Wickett *et al*., 2014) using IQ-TREE2 (Minh *et al*., 2020). Individual empirical gene alignments were concatenated using AMAS, with separate genes labeled in a partition file. The best-fit model was chosen for each partition in IQ-TREE prior to phylogeny estimation, with partitions allowed to have different evolutionary rates. Trees were inferred with 1000 bootstrap replicates. The inferred trees were transformed to ultrametric using the chronos function from the package APE (Paradis and Schliep, 2019; Paradis *et al*., 2004) in R (R Core Team, 2021) and rescaled in generations, where the number of generations was determined by dividing the total tree age by the average generation time. We determined the tree age from the original studies or the TimeTree (Kumar *et al*., 2022), while effective population size and generation time were determined from other studies (Buffalo, 2021; Cypriano-Souza *et al*., 2018; Pacifici *et al*., 2013).

The remaining 16 species trees were simulated randomly using SimPhy (Mallo *et al*., 2016), with parameter settings based on the empirical species trees properties. We set the number of taxa to 80-120 and effective population size, tree age, and generation time were varied within the limits of their values in the four empirical trees. Birth and death rates were arbitrarily set to be equal and varied between 1.96×10^*−*8^ and 4.09×10^*−*7^ depending on generation time and tree age. We visualized all simulated species trees in the ggtree package (Yu *et al*., 2017) in R, see Fig. **??**.

For each of the 20 species trees, we simulated a phylogenetic dataset of 2000 loci, with 1000 to be used for model training and validation, and the rest to be used for model testing. We relied on prior studies (Höhler *et al*., 2021, ?; Levy Karin *et al*., 2017; Willson *et al*., 2021; Zhang *et al*., 2020) to determine appropriate ranges of many simulated parameters, with other values picked arbitrarily. Overall, the parameters that varied across locus simulations included average branch length (substitution rate), variance in branch length (substitution rate variation), protein-coding (if yes, simulated under a codon model), substitution model type and parameters, locus length, Pagel’s lambda, missing data (both missing taxa and partially missing sequences), paralogy, and cross-contamination. The distribution and choice of the simulated locus properties are listed in the table **??**.

To record parameter values for each simulated locus, we adopted a simulate first approach instead of allowing tree/sequence simulators to batch simulate data. First, parameters were simulated in R. Then, SimPhy was used to simulate gene trees. A custom script then modified the trees to add deep paralogy, adjust Pagel’s lambda, and generate INDELible (Fletcher and Yang, 2009) control files. We ran INDELible on two subsets: non protein coding under NUCLEOTIDE 1 and protein coding under CODON 1. Subsequently, alignments were processed to introduce contamination, missing taxa, and missing sequence data. Raw pairwise distance and ILS levels were assessed for each dataset after simulation to confirm these higher order parameters are within reasonable ranges determined from the empirical studies (Table **??**).

### Assessment of locus properties

We selected features to assess based on prior literature (Supplemental Table 2). We assessed average UFBoot support, treeness, tree-based substitution rate mean and variance, base composition variance, saturation curve slope and correlation coefficient, taxon occupancy, alignment length, missing data percent, variable sites percent, and penalized phylogenetic informativeness (area under the curve), see table **??**.

MAFFT (Katoh and Standley, 2013) was used to align simulated and empirical loci. We used AMAS (Borowiec, 2016) to score locus length and percent of missing and variable sites. IQ-TREE2 (Minh *et al*., 2020) was used to reconstruct gene trees and a concatenated species tree, as well as determine the average UFBoot support. HyPhy (Kosakovsky Pond *et al*., 2019; Pond *et al*., 2005) was used with the inferred concatenation-based species tree to infer locus site rates needed to compute phylogenetic informativeness. Astral 5.7.8 (Sayyari and Mirarab, 2016; Zhang *et al*., 2018) was used to infer a coalescent-based species tree using gene trees (inferred in IQ-TREE and collapsed by 0 Shimodaira-Hasegawa approximate likelihood ratio test (SH-aLRT) threshold) (Guindon *et al*., 2010; Simmons and Gatesy, 2021) All other properties were computed in R using custom scripts and several packages (Dornburg *et al*., 2016; Paradis and Schliep, 2019; Paradis *et al*., 2004; Revell, Liam, ????; Schliep, 2011). For the simulated loci we also computed two proxies for locus phylogenetic utility: Robinson-Foulds (RF) similarity (1 – normalized RF distance between the inferred locus tree and true species tree) and weighted Robinson-Foulds (wRF) similarity (1 – normalized wRF). Spearman’s rho was used as a non-parametric test to determine the correlation between property values and phylogenetic utility.

To analyze the effects of high or low phylogenetic utility on difficult nodes (Fig. 2), we used the following procedure. All simulated species trees were rescaled to have a length of 1. Node heights were binned into 10 bins from 0 to 1, while branch lengths were binned into 10 bins based on log of the length, so that very short branches that are typically most difficult are more prominent in the analysis. We binned the simulated loci into three groups (RF similarity of *>*0.75, 0.25-0.75, and <0.25) and then assessed the statistical support (SH-aLRT and UFBoot) and correctness of reconstructed bipartitions across all node heights and branch length bins. We used the Wilcoxon Rank-Sum Test (Wilcoxon, 1945) with Bonferroni correction to compare the distributions of the property values with respect to RF similarity.

### Machine learning

We chose to use a Random Forest Regressor as a machine learning model to train for several reasons: the decision tree output by the Random Forest can provide insights into relative feature importance - in biological terms, the method provides distinct filtering recommendations based on locus features that allows for progressive filtration of loci to remove those with the greatest non-phylogenetic signal. Features across training subsets of all simulated datasets were combined together and split into two training datasets, one with RF similarity as the response variable, and the other with wRF (Supplementary Table 2). Each dataset had 12 features and 20000 loci.

The random forest models and were implemented and trained using the scikit-learn v1.0.1 package (Pedregosa *et al*., 2011) in python. Each training dataset was randomly split into training and testing using the default 25% of testing data. The model was initially tuned by randomized search through a parameter grid (100 iterations), followed by a more precise grid search by attempting to adjust each parameter by 25% in both directions. Parameters to tune included maximum tree depth (1000 to 20000 and no limit), maximum number of features to consider (auto, square root, and 0.5), minimum number of samples per leaf (5 to 100), number of decision trees (1000 to 10000), and minimum number of samples required for split (2 to 10). Properties of the trained models were assessed by computing mean squared error, correlation coefficient, and permutation feature importance (10 permutations). To assess the level of feature interactions in our model, we computed first-order Friedman’s H-statistic, which shows the overall level of interaction of a given feature with others, and second-order Friedman’s H-statistic, which shows the level of interactions between specific features.

### Evaluation of model-based subsampling on simulated datasets

Each of the trained random forest models (RF and wRF-based) was used to predict the phylogenetic utility of loci in the subsets of simulated loci that had not been examined by the model during testing. Due to the time that would be needed to compute inference on all subsets in all 20 simulated datasets, we limited this assessment to the four simulated datasets that were based on empirical species trees. After all loci were scored, loci within each simulation dataset were ranked by the predicted utility, and subsets of the 800 best loci (20% noise filtering), 600 best loci (40% noise filtering), and 600 worst loci, along with four randomly chosen subsets of 600 loci, were produced in order to gauge the impact of model-based filtering on the estimated phylogeny. Alignments of loci in each subset were concatenated and analyzed in IQ-TREE, while gene trees were combined and analyzed in Astral. Methodology of both types of analyses was otherwise identical to how species trees for the entire sets of loci were inferred. For the inferred IQ-TREE trees we scored RF and wRF similarity to the true species tree and average UFBoot and SH-aLRT support, while for the Astral trees we scored Robinson Foulds (RF) similarity and average local posterior probability support. Additionally, we examined the distributions of substitution rate, locus length, paralogy, and cross-contamination in the analyzed subsets. We used Wilcoxon Rank-Sum Test with Bonferroni correction to compare the distributions of the property values with respect to different subsets.

### Evaluation of model-based subsampling on empirical datasets

For each of the four empirical datasets, mentioned above, we assessed locus properties of these empirical loci similarly to the simulation datasets. We then predicted locus utility using our trained RF and (wRF) models. Each model was used to predict the phylogenetic utility of loci, and then loci were ranked based on their utility. For each dataset we then created several subsampled sets of loci, 60%, 70%, 80%, and 90% loci with the highest utility, as well as 10% loci with the highest and 10% with the lowest utility to inspect the extremes. We then analyzed the obtained subsets in concatenation, using IQ-TREE, and in coalescence framework, using Astral. We used the treespace package (Jombart *et al*., 2017) in R to obtain distances between trees and perform a principal coordinate analysis (PCoA). Wilcoxon Rank-Sum Test with Bonferroni correction was used to compare the distributions of branch length and statistical support values with respect to different subsets. Trees inferred on each subset were inspected in Figtree v1.4.3 for focal topological differences.

Visualizations were done with aid of the ggplot package (Wickham, 2016) in R. All scripts are available at https://github.com/alexandrawalling/PML

