## Supplementary Figures and Tables for "Predicting locus phylogenetic utility using machine learning"

Supplemental Figures

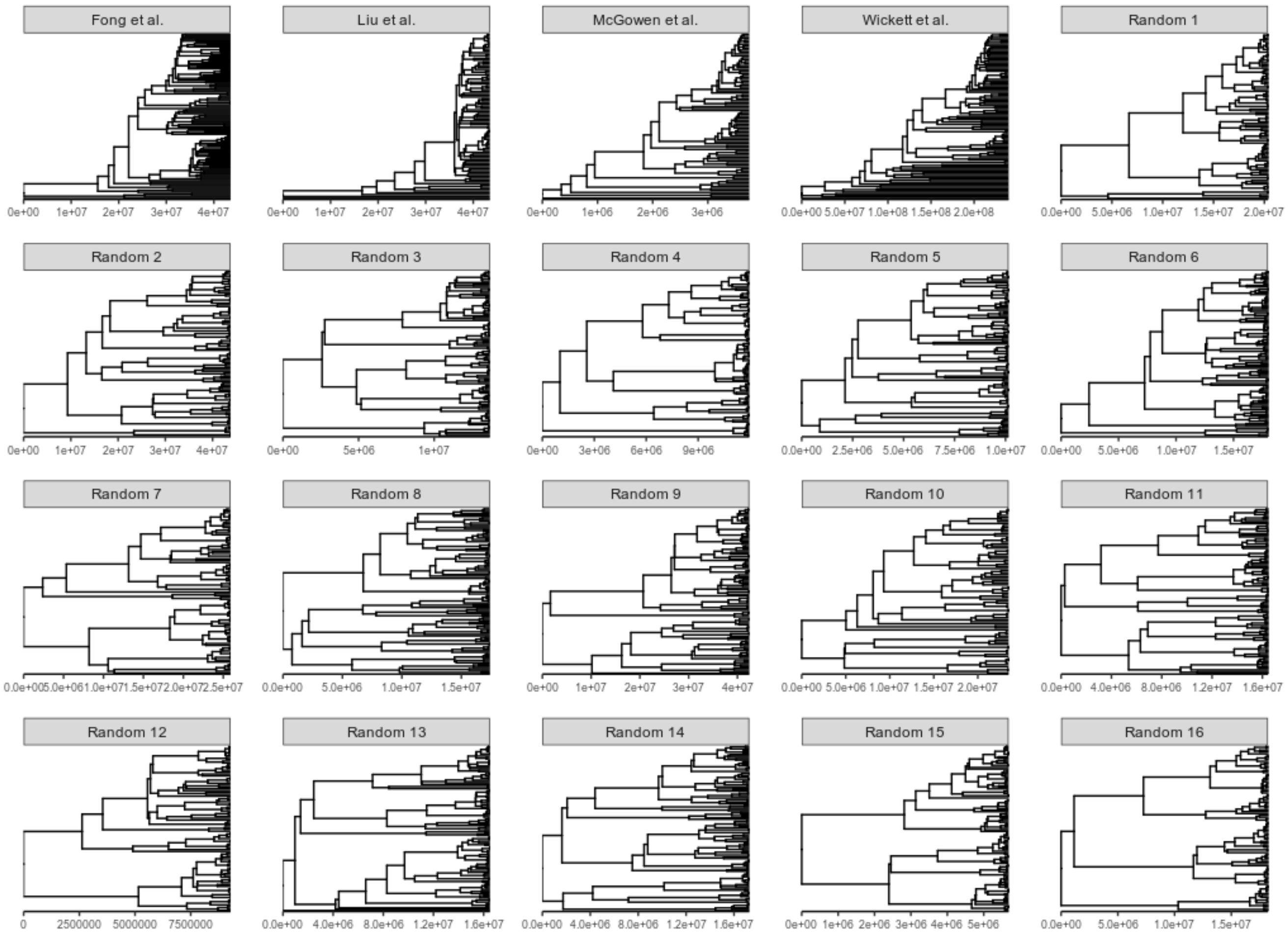


**FIG. S1.** Empirical and randomly generated species trees used for sequence dataset simulations

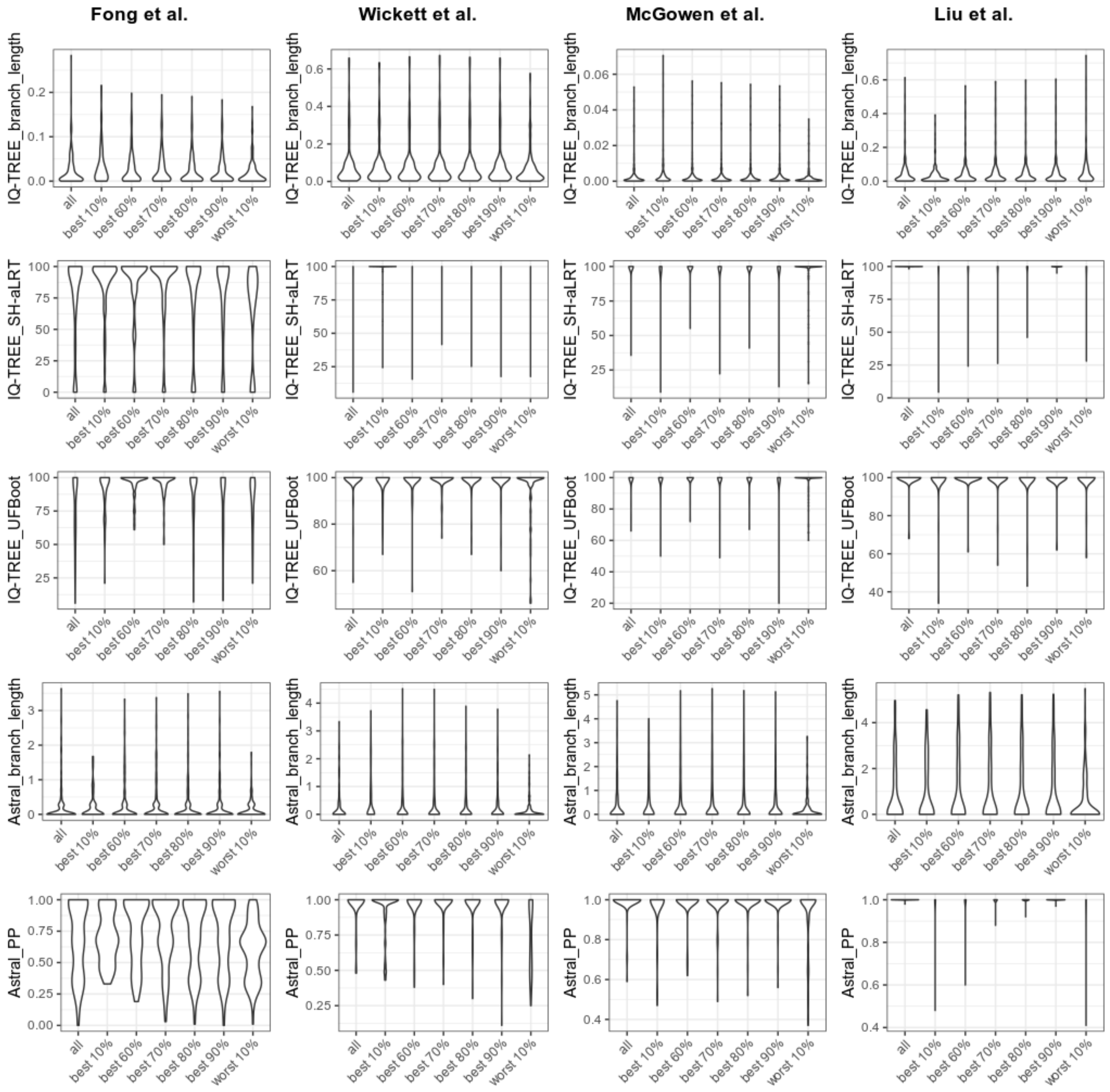


**FIG. S2.** Properties of the trees inferred on filtered empirical datasets

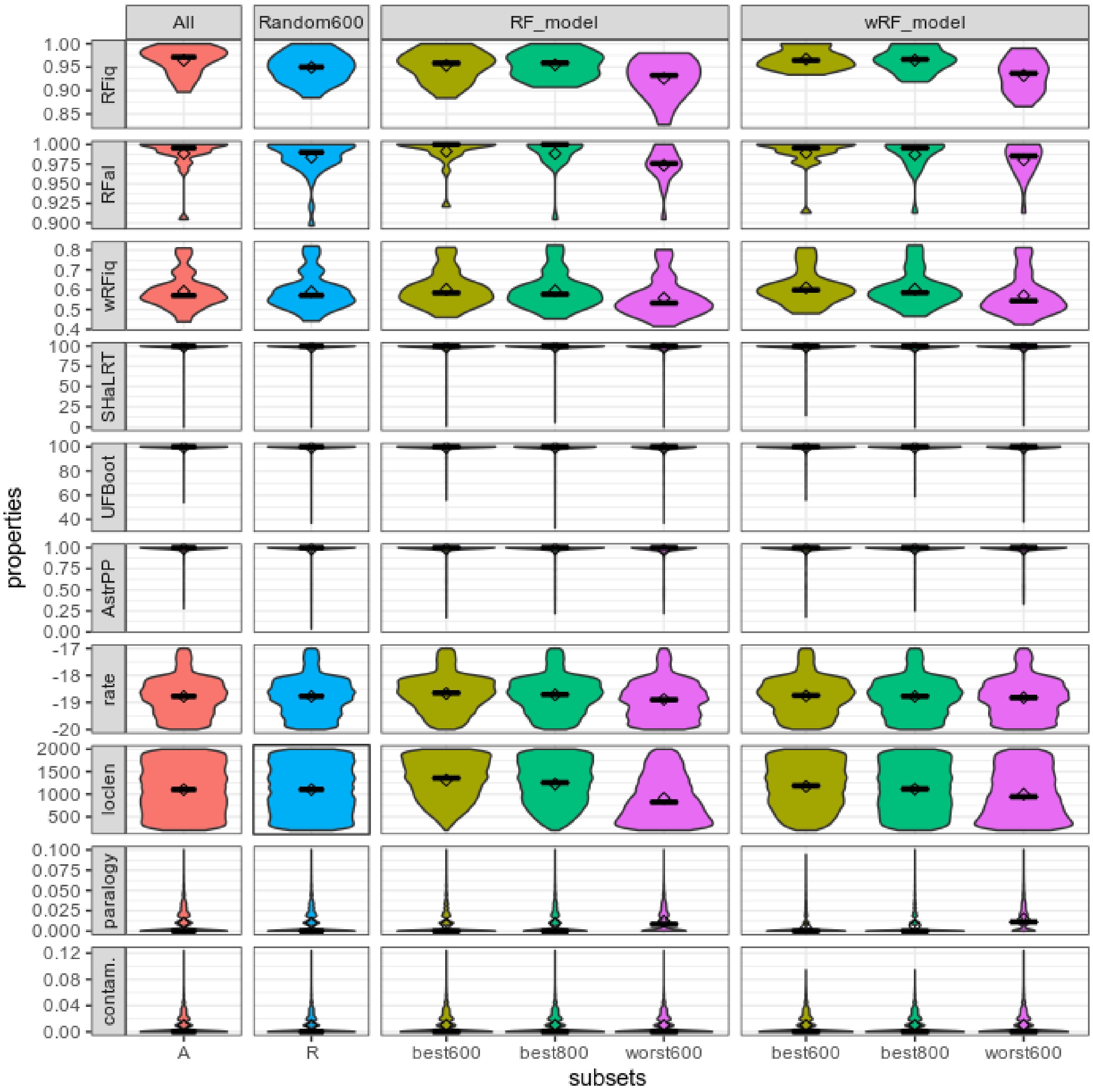


**FIG. S3.** Results of the assessment of the original and filtered sets of loci which were part of the testing dataset for the machine learning model validation. For each of the analyzed simulated datasets all 1000 loci, four random samples of 600 loci, as well as the best 800, best 600, and worst 600 loci were used to reconstruct phylogenies. Topological distance, statistical support, as well as certain properties of the subsets are shown.

Parameters used for simulation

Table S1.: Parameters used for simulation

| Property | Level | Tool | Parameters |
|---|---|---|---|
| Effective population size | Species tree | SimPhy | We randomly chose the -sp to be 1000, 10000, or 100000 for each of the species trees; these were based on the estimates for the empirical studies under consideration (see methods) |
| Taxon number | Species tree | SimPhy | We randomly chose the -sl to be between 80 and 120; this range reflected the number of taxa in the empirical studies under consideration (see methods) |
| Tree age | Species tree | SimPhy | We randomly chose the -st to be between 75 and 435 My; this range reflected the age of the groups in the empirical studies under consideration (see methods) |
| Generation time | Species tree | SimPhy | We randomly chose the -sg to be between 5 and 20; this range reflected the estimated generation time of taxa in the empirical studies under consideration (see methods) |
| Lineage birth and death rates | Species tree | SimPhy | We randomly chose the lineage rate coefficient to be between 1 and 2.5; the coefficient was divided over the ratio of tree age and generation time to obtain the resulting birth rate, which set the -sb in SimPhy; death rate was set to be equal to birth rate |

| | | | |
|---|---|---|---|
| Branch lengths, mean | Gene tree | SimPhy | We set the -su parameter to ln:X,0.1 where X was uniformly varied -20 and -18 for the random trees 1-12, -19 to -17 for the random trees 13-16, from -19.5 to -18.5 for the Fong et al. based simulated dataset, -20 to -19 for the Wickett et al. based simulated dataset, -19 to -18 for the two remaining empirical tree-based datasets. The parameters overall ranged from -20 to -17 with specific constraints to match the ranges of the pairwise distances in the empirical datasets and were consistent with the parameters chosen by some other simulation studies (Willson *et al.*, 2021; Zhang *et al.*, 2020) |
| Branch lengths, variance | Gene tree | SimPhy | we set the -hs param to ln:X,1 as in the simulator tutorial, but changed X from suggested 1.5 to vary uniformly from 0.5 to 2.5 to allow for stronger variance |
| Branch lengths, internal fraction | Gene tree | custom script | transform SimPhy generated tree using Pagel's lambda to decrease the proportion of internal branches (Pagel, 1999); value of lambda was arbitrarily chosen to be uniform 0.75 to 1.0 to match other properties of the empirical datasets |
| Substitution Model type | Gene tree | INDELible | All GTR family models (50% chance across all, each model equiprobable) and M2 codon selection model (50% chance) |
| Substitution Model change | Gene tree | INDELible | A number of model shifts was drawn according to poisson process, and applied randomly to top 25% longest branches; model shift did not involve changing protein coding, proportion of invariant (and neutral in the case of the codon model) sites, and rate heterogeneity (in the case of the nucleotide model); other parameters were drawn per model shift |

| | | | |
|---|---|---|---|
| Base Composition | Gene tree clade or taxon | INDELible | For each locus base composition was drawn from a Dirichlet distribution with alpha = 10, beta = 1, resulting in 0.25 frequency of each base, approaching the mean frequencies shown in RAxML Grove (Höhler *et al.*, 2021); For the loci under M2 model, default equal codon frequencies were retained |
| Substitution Model transition rates (GTR family models only) or kappa (M2 model) | Gene tree clade or taxon | INDELible | Truncated lognormal distributions, set based to mimic RAxML Grove results (Höhler *et al.*, 2021) |
| Substitution Model rate heterogeneity (GTR family models only) | Gene tree | INDELible | we used continuous gamma distribution for 50% of loci, alpha was set based on RAxML Grove results (Höhler *et al.*, 2021) |
| Substitution Model indel model | Gene tree clade or taxon | INDELible | sampled from uniform distribution from 1.5 to 2.0, consistent with results of Levy Karin *et al.* (2017) |
| Substitution Model indel rate | Gene tree clade or taxon | INDELible | sampled from uniform distribution from 0.001 to 0.002, consistent with results of Levy Karin *et al.* (2017) |
| Substitution Model proportion of sites (excluding invariant) under selection (M2 model only) | Gene tree | INDELible | We chose a uniform distribution from 0 to 1, with the remainder of sites being under strictly neutral selection |
| Substitution Model selection strength (M2 model only) | Gene tree clade or taxon | INDELible | We arbitrarily chose a uniform distribution from 0 to 3 to obtain a varying degree of selection ranging from purifying to strong positive |

| | | | |
|---|---|---|---|
| Substitution Model proportion of invariant sites | Gene tree | INDELible | Since empirical proportion of invariant sites may be correlated with rate, we instead treated this parameter as fraction of sites under selection to remain unchanged, and sampled it from a uniform distribution, arbitrarily set from 0 to 0.25 |
| Alignment length | Gene tree | INDELible | sampled from uniform 200-2000bp as this range encompasses mean lengths of loci in many modern phylogenomic datasets, including the empirical datasets chosen in this study |
| Missing taxa | Gene tree | custom script | random 0 to 50% of taxa selected and removed as it is customary to filter out alignments with less 50% of taxa (Dietrich *et al.*, 2017; Lemmon *et al.*, 2009; Young *et al.*, 2016) |
| Missing segments | Gene tree | custom script | among the remaining taxa, random up to 50% taxa selected, for each random 20-60% missing amount is drawn; to simplify the procedure, we confined the remaining data present to be a single continuous present, thus missing data was confined to one or both flanks; the balance of missing data between left and right flanks was determined by a flank bias parameter uniformly varied between 0 and 1; flanking missing data is more common as it coincides with low coverage introns flanking the chosen exonic loci or flanking poorly enriched regions in the hybrid enrichment experiments |

| | | | |
|---|---|---|---|
| Deep paralogy | Gene tree | custom script | Deep paralogy modeled here is associated with spurious undesired sequences remaining in loci due to processing errors, and will likely vary depending on a dataset and a pipeline; we thus arbitrarily specified a zero inflated poisson 0.5 distribution to model both presence and number of paralogs per locus and 50% of loci be paralogy-free; paralogous taxa for a locus were randomly determined modified to form a separate clade sister to the rest of taxa on a branch which length was sampled uniformly to be from 1 to 10 times longer than the longest branch of the original gene tree |
| Contamination | Gene tree | custom script | The distribution of contaminant pairs presence and number was modeled similar to deep paralogy; for each count a pair of taxa was randomly determined and sequence of one of the taxa was copied in the other taxon |

Assessed Properties

Table S2.: Assessed locus properties

| Property | Assessment methods | References |
|---|---|---|
| Base composition variance | computed frequency of each base (ATCG) in each sequence (taxon), computed variance of each base frequency across all taxa, summed up the variances | Evangelista *et al.* (2021); Kocot *et al.* (2017); Mongiardino Koch and Thompson (2021); Whelan *et al.* (2015) and others |
| Alignment length | alignment length computed by AMAS (Borowiec, 2016) | Brown and Thomson (2017); Evangelista *et al.* (2021); Gernandt *et al.* (2018); Mongiardino Koch and Thompson (2021) and others |
| Proportion of missing data | percent missing computed by AMAS (Borowiec, 2016) | Brown and Thomson (2017); Evangelista *et al.* (2021); Herrando-Moraira and Cardueae Radiations Group (2018); Kocot *et al.* (2017); Molloy and Warnow (2018); Mongiardino Koch and Thompson (2021) and others |
| Taxon occupancy | ratio btw N of terminals of gene tree and species tree | Borowiec *et al.* (2015); Dietrich *et al.* (2017); Evangelista *et al.* (2021); Gernandt *et al.* (2018); Herrando-Moraira and Cardueae Radiations Group (2018); Lemmon *et al.* (2009); Mongiardino Koch and Thompson (2021); Young *et al.* (2016) and others |
| Proportion of variable sites | proportion of variable sites computed by AMAS (Borowiec, 2016) | Mongiardino Koch and Thompson (2021) and others |

| | | |
|---|---|---|
| Substitution rate mean and variation | Tree-based mean rate was computed as described by (Mongiardino Koch and Thompson, 2021), using the inferred gene trees; rate variation was computed simply a variance of all gene tree branch lengths | Borowiec *et al.* (2015); Gernandt *et al.* (2018); Herrando-Moraira and Cardueae Radiations Group (2018); Mongiardino Koch and Thompson (2021); Whelan *et al.* (2015) and others |
| Treeness | calculated as in (Mongiardino Koch and Thompson, 2021) | Mongiardino Koch and Thompson (2021) |
| Saturation | saturation regression line coefficient and R-squared as in Borowiec | Borowiec *et al.* (2015); Herrando-Moraira and Cardueae Radiations Group (2018); Kocot *et al.* (2017); Mongiardino Koch and Thompson (2021) and others |
| Average support | mean value of UFBoot | Borowiec *et al.* (2015); Herrando-Moraira and Cardueae Radiations Group (2018); Mongiardino Koch and Thompson (2021) and others |
| Phylogenetic Informativeness | Penalized Phylogenetic Informativeness, area under the curve, as in Mongiardino Koch and Thompson (2021) | Herrando-Moraira and Cardueae Radiations Group (2018); Mongiardino Koch and Thompson (2021) |

Table S3.: Raw pairwise distance and ILS of each dataset

| Dataset | RPD, mean | RPD, range | ILS, mean | ILS, range |
|---|---|---|---|---|
| Fong et al. (empirical) | 14.016% | 3.659-24.400% | N/A | N/A |
| Wickett et al. (empirical) | 28.35% | 8.40-50.65% | N/A | N/A |
| McGowen et al. (empirical) | 2.4687% | 0.2622-7.0602% | N/A | N/A |
| Liu et al. (empirical) | 15.97% | 5.62-33.53% | N/A | N/A |
| Simulated on Fong et al. tree | 13.180% | 1.575-42.711% | 6.252% | 1.587-10.317% |
| Simulated on Wickett et al. tree | 29.338% | 4.571-60.476% | 0.001% | 0.000-1.000% |
| Simulated on McGowen et al. tree | 2.7592% | 0.3163-13.0841% | 0.000% | 0.000-0.000% |
| Simulated on Liu et al. tree | 13.724% | 2.574-42.556% | 0.001149% | 0.000000-1.149425% |
| Simulated tree 1 | 8.191% | 1.034-34.217% | 8.144% | 1.111-16.667% |
| Simulated tree 2 | 16.852% | 2.944-53.776% | 3.448% | 0.000-8.696% |
| Simulated tree 3 | 7.243% | 0.676-30.419% | 0.6681% | 0.0000-3.9216% |
| Simulated tree 4 | 6.5606% | 0.6875-27.7679% | 0.03654% | 0.00000-0.96154% |
| Simulated tree 5 | 5.9487% | 0.6252-26.0544% | 0.000% | 0.000-0.000% |
| Simulated tree 6 | 7.956% | 1.124-33.227% | 7.584% | 1.905-15.238% |
| Simulated tree 7 | 12.224% | 1.225-43.660% | 0.04713% | 0.00000-1.14943% |
| Simulated tree 8 | 9.801% | 1.352-36.738% | 0.03843% | 0.00000-1.85185% |
| Simulated tree 9 | 16.557% | 2.696-56.059% | 0.702% | 0.000-2.000% |
| Simulated tree 10 | 11.509% | 1.902-41.255% | 0.1769% | 0.0000-1.2821% |
| Simulated tree 11 | 9.233% | 1.425-35.570% | 0.8597% | 0.0000-4.0816% |
| Simulated tree 12 | 5.4145% | 0.4549-25.0981% | 11.057% | 1.149-20.690% |
| Simulated tree 13 | 17.850% | 3.713-55.472% | 9.123% | 1.754-16.667% |
| Simulated tree 14 | 18.929% | 3.452-59.218% | 5.197% | 0.000-11.304% |
| Simulated tree 15 | 7.775% | 1.003-29.659% | 25.43% | 13.21-39.62% |

| | | | | |
|---|---|---|---|---|
| Simulated tree 16 | 18.857% | 1.287-56.940% | 1.629% | 0.000-5.814% |
